# The effect of genetic architecture and selfing on the capacity of a population to expand its range

**DOI:** 10.1101/2020.01.22.915454

**Authors:** Martin Eriksson, Marina Rafajlović

**Affiliations:** Department of Marine Sciences, University of Gothenburg, Gothenburg, Sweden; The Linneaus Centre for Marine Evolutionary Biology, University of Gothenburg, Gothenburg, Sweden; Gothenburg Global Biodiversity Centre, University of Gothenburg, Gothenburg, Sweden

**Keywords:** Recombination, range margins, range contraction, genetic variation, adaptation, simulations

## Abstract

Previous theoretical work on range expansions over heterogeneous environments showed that there is a critical environmental gradient where range expansion stops. For populations with freely recombining loci underlying the trait under selection (hereafter *adaptive loci*), the critical gradient in one-dimensional habitats depends on the fitness cost of dispersal, and the strength of selection relative to genetic drift. Here, we extend the previous work in two directions and ask: What is the role of the genetic architecture of adaptive loci during range expansions? And what effect does the ability of selfing have on range expansions? To answer these questions, we use computer simulations. We demonstrate that, while reduced recombination rates between adaptive loci slow down range expansions due to poor purging of locally deleterious alleles at the expansion front, they may also allow a species to occupy a greater range. In fact, for some parameter values, we find that a population with freely recombining adaptive loci experiences global extinction, whereas reduced recombination rates allow for a successful expansion over a wide geographic range. In addition, we find that allowance of selfing may improve the ability of populations to expand their ranges. We discuss the mechanisms underlying these results.

## Introduction

As environmental conditions have changed in the past, ranges of many populations have shifted, contracted or expanded. Occasionally, range expansions and shifts have resulted in the creation of new species (Abbot and Double 2003; Bergström et al. 2005; Van Bocxlaer et al. 2010; Momigliano et al. 2018). In some cases, by contrast, species or their local populations have gone extinct because they have failed to adapt to the new conditions or failed to change their ranges accordingly (Benton and Twitchett 2003). Today, human activities, such as environmental degradation, introduction of foreign species and anthropogenic climate change, put the current ecosystems under unprecedented stress (Lewis and Maslin 2015; Waters et al. 2016). Hence, it is expected that many species will have to change their ranges in the future to avoid extinction (Davis and Shaw 2001; Hill et al. 2011; Urban 2015).

One of many factors that can limit a species’ range is that asymmetric gene flow from the more densely populated source populations can preclude adaptation to the sparsely populated front of the expansion, by swamping of locally adapted alleles (Mayr 1963; Peck et al. 1998). When the environmental gradient is nonlinear, theoretical models, supported by recent empirical evidence (Bachmann et al. 2019), have shown that a population manages to adapt and colonise the habitat up to the point where the steepness of the gradient reaches a critical threshold, and here a stable range margin forms (Polechová and Barton 2015; Polechová 2018; Bridle et al. 2019). For finite populations, this critical threshold occurs at a steepness where the genetic load, generated by gene flow, reduces the mean growth rate of local populations at the expansion front to the point where genetic drift causes adaptation to fail (Polechová and Barton 2015; Polechová 2018). Recently, it has been demonstrated that stable range margins can also form when environmental gradients are interrupted by a flat segment (Bridle et al. 2019).

Importantly, the studies that found stable range margins due to steepening or interrupted environmental gradients (e.g. Bridle et al. 2010; Polechová and Barton 2015; Polechová 2018; Bridle et al. 2019) were assuming free recombination between the loci underlying the trait under selection (hereafter *adaptive loci*). However, the genome-wide recombination rate together with the gene density are highly variable between populations (Mezard 2006; Ellegren and Galtier 2016; Ravinet et al. 2017; Stapley et al. 2017). Due to its two antagonistic effects (Felsenstein 1974; and see below), the recombination rate between adaptive loci may be a key component of local adaptation and, consequently, range expansion. To date, however, the role of recombination during range expansions has not been elucidated.

The effect of recombination is nontrivial, as pointed out already by Felsenstein (1974). More frequent recombination is more effective in purging deleterious alleles from a population, as well as in bringing together beneficial alleles segregating in different individuals (Felsenstein 1974). During range expansions, effective purging mechanisms might be particularly important due to the so-called gene-surfing effect wherein potentially locally deleterious alleles may increase in frequency and even fix at the expansion front where populations are small (Hallatschek and Nelson 2008; Slatkin and Excoffier 2012). This effect of genetic drift, known as *expansion load*, may lead to a reduction in local fitness at the expansion front (Peischl et al. 2013). Note that a consequence of a higher expansion load is a slower expansion, both over spatially homogeneous (Peischl et al. 2015) and over heterogeneous environments (Gilbert et al. 2017). In the former case (spatially homogeneous environments) it was shown that expansion load is more severe in the absence than in the presence of recombination between adaptive loci (Peischl et al. 2015).

By contrast, for range expansions over more realistic scenarios - where stable range margins are expected to form (e.g. with a steepening gradient) - the effect of recombination has not been investigated to date. However, and somewhat counter-intuitively, Gilbert et al. (2017) showed that, when expansion occurs over spatially heterogeneous environments, expansion load is smaller than when environmental conditions are spatially homogeneous. This is due to dispersal between local populations subject to different environmental conditions: it primarily slows down expansion and, at the same time, reduces the effect of genetic drift at the expansion front, making selection more efficient (Gilbert et al. 2017).

While Gilbert et al. (2017) did not investigate how different recombination rates between adaptive loci may influence expansion load and the resulting dynamics of range expansion, their finding suggests that an increase in expansion load due to reduced recombination *sensu* Peischl et al. (2015) (in a model with constant environmental conditions and absence versus presence of recombination) may be counteracted by the effect of local maladaptation that, instead, reduces expansion load in heterogeneous environments.

In addition to the above, less frequent recombination between adaptive loci (tight genetic architectures) may, in fact, be advantageous over frequent recombination (loose genetic architectures). This is because blocks of locally well adapted alleles are more effectively protected when they are harboured by a region of suppressed recombination, e.g. a colinear genomic region with tightly linked loci, or a chromosomal inversion (Kirkpatrick and Barton 2006; Kirkpatrick and Barrett 2015). This effect may, thus, additionally counteract the above mentioned effect that reduced recombination increases expansion load (*albeit* in a model with constant environmental conditions; Peischl et al. 2015).

Another factor that may be relevant during range expansions, besides the recombination rate, is the mode of reproduction. Previous studies on range expansion focused either on clonal reproduction (Peischl et al. 2015), or on fully sexual populations (Polechová and Barton 2015; Polechová 2018; Bridle et al. 2019). Here we ask how the ability to self-fertilise affects the capability of a population to expand its range compared to when selfing is not possible. Selfing is common among plants and some hermaphrodite animals (Jarne and Charlesworth 1993), and it is known that uniparental reproduction can increase the speed of range expansion, at least under spatially homogeneous conditions (Baker 1955; Tomlinson 1966; Pannell and Barrett 1998; Rafajlović et al. 2017). However, the ability for a single selfing individual to establish a new colony might come at a price of locally reduced genetic variation. This may reduce the ability of a population to further expand its range over a heterogeneous environment. This suggests that both the effect of allowance or disallowance of selfing and the effect of tight versus loose genetic architectures on range expansions over spatially heterogeneous environments are nontrivial, but neither have been investigated so far.

Here we study the interplay between expansion load, local maladaptation due to dispersal in spatially heterogeneous environments and genetic architecture underlying the trait under selection (more or less tightly linked adaptive loci) in populations with and without selfing. Obtaining this knowledge is crucial to understand the mechanisms by which natural populations may extend their ranges, and to which extent. More specifically, we ask: What is the role of the genetic architecture of adaptive loci and the mode of reproduction (selfing allowed or not) during range expansions? Do tighter genetic architectures in populations either with or without selfing alter the ability of a population to expand its range in terms of the speed of the expansion and the critical gradient? Do the answers to these questions depend on the amount of standing genetic variation in the source population? If yes, how?

To address these questions, we performed individual-based simulations, extending the model proposed by Polechová and Barton (2015) to incorporate arbitrary recombination rates between adaptive loci, to vary the amount of standing genetic variation in the source population, and to allow or preclude selfing. As expected, we found that range expansion is slower when genetic architecture is tighter. However, we also found that, for many parameter values in our model, tighter genetic architectures (but not too tight) and/or selfing allow populations to expand their ranges over steeper environmental gradients than those attained with freely recombining adaptive loci. This effect is transient, but long-lasting with persistence time of tens of thousands of generations. This time period is specifically important for populations that have expanded their ranges after the last glacial period (approximately during the last 10, 000 years; Clark et al. 2009; Andrén et al. 2011). Importantly, we found that reduced recombination allows the population to more efficiently utilise genetic variation: for some parameter values, we found that a population with high standing genetic variation can expand its range only with tight genetic architectures, whereas it experiences global extinction otherwise. In conclusion, there is a trade-off between short-term speed of range expansion and the range that a population may attain and pursue on the long run.

## Methods

We used individual-based simulations to investigate how different recombination rates between adaptive loci in populations with and without selfing affect range expansion. The habitat under consideration was modelled as a one-dimensional chain of *M* demes of equal sizes (notations for parameters are summarised in table 1). We measured distances between demes in units of the number of demes. The local carrying capacity in each deme was set to *K* = 150. We assumed an environmental gradient along the habitat, similarly to Polechová and Barton (2015) and Bridle et al. (2019), such that the optimal phenotype in deme *i* = 1, …, *M*, denoted by *θ*_*i*_, was assumed to be a cubic polynomial of *i*, with an inflection point at the center of the habitat (see Appendix B for details).

**Table 1:** Explanation of the parameters of the model, and the notations used throughout.

| Parameter | Description |
| --- | --- |
| $M$ | Number of demes in the habitat |
| $K$ | Local carrying capacity |
| $\theta_i$ | Optimal phenotype in deme $i$ |
| $L$ | Number of loci under selection |
| $\alpha$ | Allele-effect size |
| $r_m$ | Maximal intrinsic growth rate |
| $\mu$ | Mutation rate |
| $V_S$ | Width of stabilising selection |
| $\sigma$ | Standard deviation of Gaussian dispersal function |
| $c$ | Recombination rate |

Generations were discrete and non-overlapping. Populations were diploid and monoecious. We assumed that *L* bi-allelic loci underlain the trait under selection. At each locus under selection, we further assumed that alleles had two possible allele-effect sizes, *α* > 0 or 0. The phenotype of each individual was equal to the sum of allele-effect sizes at the *L* loci under selection (Table 1).

The life-cycle was: selection → recombination → mutation of gametes → fertilisation → migration. Thus, migration occurs at the seed or larval stage, as for most selfing species, such as plants or sessile invertebrates. The fitness of each individual was assumed to depend on the individual’s phenotype and on the local population density. Similarly to Polechová and Barton (2015), but assuming a diploid population, the number of gametes, 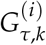, produced in generation *τ* in deme *i* by individual *k* was drawn from the Poisson distribution with mean 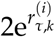 where

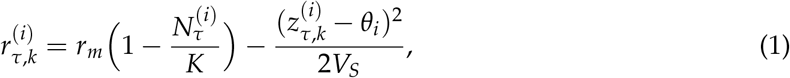

is the corresponding growth rate. In eq. (1), *r*_*m*_ denotes the maximal intrinsic growth rate, *V*_*S*_ denotes the width of stabilising selection, 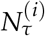 is the number of individuals in deme *i* in generation *τ*, and 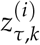 is the phenotype of individual *k* in deme *i* in generation *τ*. As in Polechová and Barton (2015), we assumed that *r*_*m*_ and *V*_*S*_ are the same for all individuals, in all demes, and in all generations. In the model, recombination occurred with probability *c* between each pair of adjacent loci. After recombination, one of the homologous sets of alleles was chosen at random to generate a gamete. In generation *τ*, this process was repeated 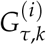 times for individual 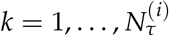 in deme *i* = 1, …, *M*. With this, a pool of 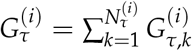 gametes was created in deme *i* for the next generation.

Mutations occurred symmetrically, so that an allele with effect size *α* was mutated to an allele with effect size 0, and vice versa, with equal probabilities, either at rate *µ* = 10^*−*8^ or at rate *µ* = 10^*−*6^ per gamete per locus per generation. During the fertilisation phase, a pair of two gametes from the same deme was chosen at random, without replacement, until the pool of gametes was exhausted or until there was only a single gamete left (which was discarded). For each set of parameters, one set of simulations was performed with selfing allowed at no cost, and another set with selfing disallowed. When selfing was allowed at no cost, the rate of selfing was not fixed in the model, but depended on the number of individuals and their relative fitnesses (see below). In the idealised case, when all individuals have the same fitness, the rate is 1/*N*. Note that when *N* grows over time, the realised rate of selfing decreases. When selfing was not allowed, the offspring died if both gametes came from the same parent. Note that, in the model by Polechová and Barton (2015), the local population was eliminated in each deme that had less than 5 individuals in a given generation, but here we did not make use of this constraint. Finally, migration was assumed to occur according to a Gaussian function with mean 0, and standard deviation denoted by *σ* (see Appendix B for details).

At the start of each simulation, the population was assumed to occupy a fraction of the habitat, i.e. *M*/5 demes arranged side-by-side around the centre of the habitat, with exactly *K* individuals in each occupied deme. All other demes were assumed to be empty. The starting mean phenotype in a locally occupied deme was assumed to be equal to the environmental optimum in that deme. Initially, the adaptive loci were assumed to be in linkage equilibrium. To generate the starting genotypes, 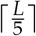 loci (where 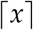 denotes the smallest integer larger than or equal to *x*) were first selected at random, and assigned allele frequencies according to the clines predicted by Barton (2001), so that the trait mean in each initially occupied deme matched the optimum in that deme. The allele frequencies at the remaining loci were chosen in such a way that the trait mean of the population was left unchanged. This was achieved either by choosing uniformly at random half of the remaining loci homozygous for alleles with effect size *α* and the other half homozygous for alleles with effect size 0, or by letting the alleles at all the remaining loci take the effect size *α* with probability 0.5, and otherwise take the effect size 0. The former setting corresponded to a population with low genetic variation at the start of the expansion, i.e. the equilibrium for a population consisting of *M*/5 demes in the absence of mutations (Barton 2001). The extent of genetic variation in the latter setting was maximised, such that the range expansion was driven by older standing genetic variation rather than by solely novel mutations. Alternatively, such high genetic variation may be attained shortly after secondary contact between populations well adapted to the conditions in the left and right *M*/10 demes around the centre of the habitat (assuming that homozyguous loci were fixed for the alternative alleles at the opposite sides of the habitat centre before the contact).

The sets of parameter values considered in each simulation are summarised in table 2. Because most parameters relevant to range margins have been extensively studied by others (e.g. Polechová and Barton 2015; Bridle et al. 2019), our main focus was to investigate how different genetic architectures (i.e. different recombination rates between adaptive loci), together with selfing affected range expansion, while keeping other parameters fixed. In simulations where the initial standing genetic variation was high, the mutation rate was set to *µ* = 10^*−*8^, which is close to the human mutation rate (Nachman and Crowell 2000). Conversely, in simulations where the initial standing genetic variation was low, the mutation rate was set to *µ* = 10^*−*6^ which allowed us to increase the speed of the simulations. However, to check for the effect of the mutation rate in simulations with low initial genetic variation, we also used *µ* = 10^*−*8^ in some cases. The allele-effect size *α* and the width of stabilising selection were chosen within the range of parameter values considered by Polechová and Barton (2015), and were either set to 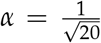, *V*_*S*_ = 2 or 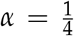, 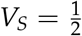, corresponding to weak or strong selection per locus, respectively. The maximal intrinsic growth rate and the dispersal parameter were fixed to *r*_*m*_ = 1.025 and 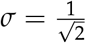, respectively, both moderately high values used by Polechová and Barton (2015). We used recombination rates of *c* = 10^*−*5^, 10^*−*4^, 10^*−*3^, 5 · 10^*−*3^, 0.01, 0.05 and 0.5.

**Table 2:** Sets of parameter values used in the simulations.

| Parameter | Low initial standing genetic variation |  | High initial standing genetic variation |  |
| --- | --- | --- | --- | --- |
|  | Parameter set LW | Parameter set LS | Parameter set HW | Parameter set HS |
| L | 574 | 513 | 574 | 513 |
| $\alpha$ | $\frac{1}{\sqrt{20}}$ | $\frac{1}{4}$ | $\frac{1}{\sqrt{20}}$ | $\frac{1}{4}$ |
| $V_S$ | 2 | $\frac{1}{2}$ | 2 | $\frac{1}{2}$ |
| $\mu$ | $10^{-6}$ | $10^{-6}$ | $10^{-8}$ | $10^{-8}$ |

Note: Abbreviations (LW, LS, HW, HS): L (H) stands for low (high) standing genetic variation, and W (S) stands for weak (strong) selection. In all cases, the number of demes was $M = 180$ , the local carrying capacity was $K = 150$ , the maximal intrinsic growth rate was $r_m = 1.025$ and the standard deviation of the Gaussian dispersal function was $\sigma = \frac{1}{\sqrt{2}}$ . Simulations for each parameter set were run both with selfing disallowed and with selfing allowed at no cost, and with recombination rates $c = 10^{-5}, 10^{-4}, 10^{-3}, 5 \cdot 10^{-3}, 0.01, 0.05$ and $0.5$ . Low genetic variation in this case means that $V_G = 0.13$ or $V_G = 0.05$ in the central deme of the habitat for weak and strong selection, respectively. High genetic variation means that $V_G = 11.6$ or $V_G = 12.9$ in the central deme of the habitat for weak and strong selection, respectively.

The number of demes in the habitat was chosen in such a way that the habitat was at least 10 times wider than the expected cline width in the case of freely recombining loci, as per the prediction by Barton (2001) (see Appendix B, where we also list the corresponding cline widths obtained under our model, calculated using the formula from Polechová and Barton 2011, p. 228). Here, we used *M* = 180. The number of loci was chosen in such a way that if all loci were homozygous for alleles of effect size *α*, then the phenotype would equal the optimal phenotype in the deme *M*, i.e. *θ_M_* = 2*Lα*. For our parameter values, the number of loci was *L* = 574 for 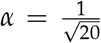 and *L* = 513 for 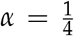.

In one-dimensional habitats (assuming that *c* = 0.5), it has been predicted that, when an environmental gradient becomes sufficiently steep, adaptation will eventually fail due to positive feedback between maladaptive gene flow and genetic drift (Polechová and Barton 2015), and stable range margins will form. The range margins of a population are determined by two composite dimensionless parameters, the *effective environmental gradient*, 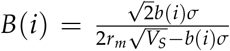 (where 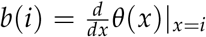 is the slope of the phenotypic optimum), and the strength of selection relative to drift, 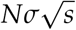 (where 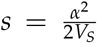 is the strength of selection per locus). When 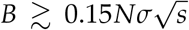, adaptation fails (Polechová and Barton (2015)). In our model, this means that for the weak selection parameter set (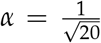, *L* = 574, *V*_*S*_ = 2) the adaptation will fail in regions that are more than 65 demes away from the centre of the habitat in each direction. For the strong selection parameter set (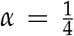, *L* = 513, 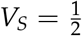) the adaptation will fail in regions that are more than 50 demes away from the centre of the habitat in each direction. When range expansion occurs over a gradually steepening gradient, as in our case, it is expected that a few additional demes will be occupied by migrants from the regions with shallower environmental gradients (cf. the results by Polechová and Barton 2015). When our simulations were initialised with low standing genetic variation, and were run without selfing and with free recombination (see Results below), we found that approximately 136 demes were occupied for the weak selection parameter set and approximately 102 demes were occupied for the strong selection parameter set, when the steady state was obtained. These ranges are hereafter referred to as the *critical range* for weak and strong selection, respectively.

For each set of parameters, 50 independent realisations of the model were performed. The local genetic variation and local population size were recorded every 500 generations. The local genetic variation was measured using the genetic variance (*V*_G_) defined as (Barton 2001)

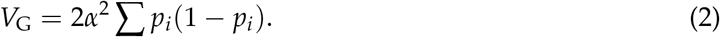

Here, *p*_*i*_ denotes the frequency of alleles with allele-effect size *α* in deme *i*. A genetic variance greater than 2.5 indicates that about one fifth of the adaptive loci are polymorphic. Conversely, the maximum value of *V*_G_ is obtained when all individuals are heterozygotes at all adaptive loci. For our parameter values corresponding to weak, and strong selection (see above), the maximum value of *V*_G_ equals 14.35, and 16.03, respectively. Given a value of *V*_G_, the average growth rate of the entire population in deme *i* and at generation *τ* is given by

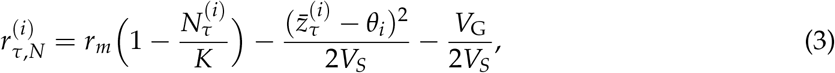

where 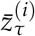 denotes the average phenotype in deme *i* at generation *τ*.

We measured the capability of a population to expand its range using the *total expansion range* (see next), and the effective environmental gradient at the expansion front. We considered demes that contained 5 or more individuals to be occupied. Because in some cases the occupied demes may be mutually disconnected (i.e. intermingled by unoccupied demes), we measured the *total expansion range* as the distance (in units of the number of demes) between the last occupied demes at the two edges of the occupied range. The total expansion range was measured shortly after the start of the expansion (i.e. after 500 generations), at its maximum and also 200, 000 generations after the start of the expansion. The significance of differences between the total expansion range obtained in simulations with different parameter values were analysed using the Wilcoxon-Mann-Whitney test (Mann and Whitney 1947).

The computer codes used in this study are available from the authors upon request.

## Results

The speed of range expansion decreased with decreasing recombination rate (*c*) below 5 ⋅ 10^*−*3^. For *c* ≥ 5 ⋅ 10^*−*3^, there was no significant difference in the speed of expansion for different values of *c* (figs. 1–2, fig. A1). The effect of different recombination rates on the speed of expansion also depended on the selection strength per locus, and the amount of initial (standing) genetic variation in the source population. When selection was weak, populations with high standing genetic variation expanded faster than populations with low standing variation, except in the case when the recombination rate was too low (*c* = 10^*−*5^; compare the corresponding coloured lines in fig. 1A and C when selfing was not allowed; as well as in fig. 1B and D for the model with allowed selfing).

**Figure 1:**
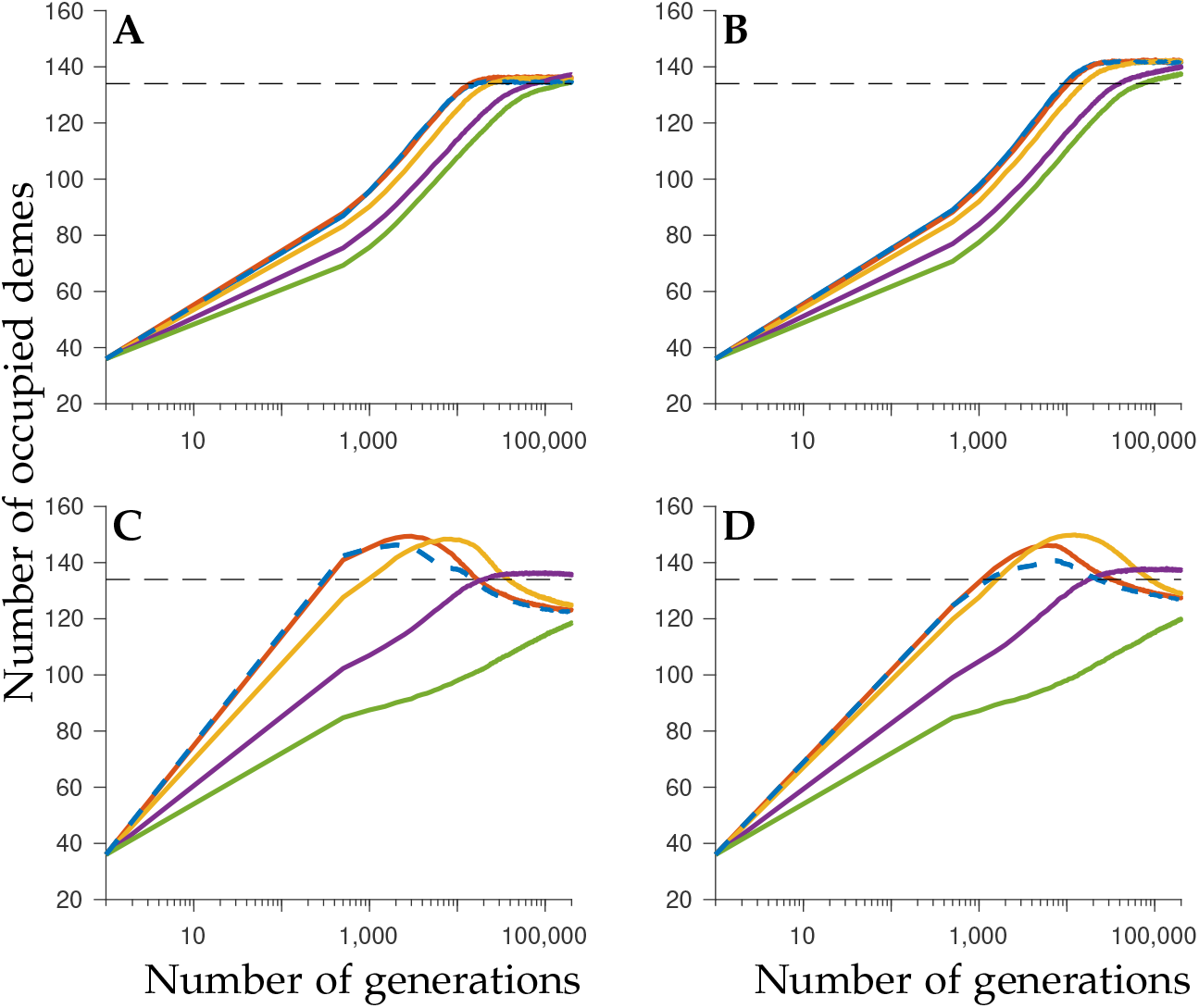
Temporal evolution of the mean number of occupied demes for weak selection. The results correspond to the model with low standing genetic variation (parameter set LW; A, B) or high standing genetic variation (parameter set HW; C, D). Selfing is either not allowed (A, C) or it is allowed at no cost (B, D). Dashed lines show the expected number of occupied demes in the steady state. Different colours denote the results with different recombination rates: *c* = 0.5 (i.e. free recombination; dashed blue), *c* = 5 ⋅ 10^*−*3^ (red), *c* = 10^*−*3^ (yellow), *c* = 10^*−*4^ (purple), *c* = 10^*−*5^ (green).

By contrast, when selection was strong (fig. 2), the joint effect of recombination rate, selfing and standing genetic variation on range expansion was more complex than in the case of weak selection. When selfing was not allowed, the entire population went extinct within a few generations when *c* ≥ 5 ⋅ 10^*−*3^ and the simulations were initialised with high standing genetic variation (fig. 2C). Thus, the allowance of selfing (fig. 2D), and/or a recombination rate of *c* ≤ 10^*−*3^ between the adaptive loci when selfing was not allowed (fig. 2C, yellow, purple, green) rescued the population from extinction (although 7 out of 50 realisations led to extinction also when *c* = 10^*−*3^ and selfing was not allowed). In the corresponding model with low standing genetic variation, however, the population expanded its range successfully for all values of *c*, either with or without selfing (fig. 2A, B). In the cases when global extinction did not occur, we observed that the speed of expansion was smaller when the recombination rate between the adaptive loci was smaller, similarly as when selection was weak. Note that for the simulations initialised with high standing genetic variation, the initial genetic variance was *V*_G_ = 11.6 or *V*_G_ = 12.9 in the central deme of the habitat for weak and strong selection, respectively (table 2). From equation (3), it follows that the initial average growth rate of the population was thus 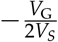 in the center of the habitat (where *N* = *K* and 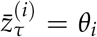). This means that the initial average growth rate for the population was 2.9 for weak selection and 12.9 for strong selection, in the centre of the habitat, when the simulations were initialised with high standing genetic variation.

**Figure 2:**
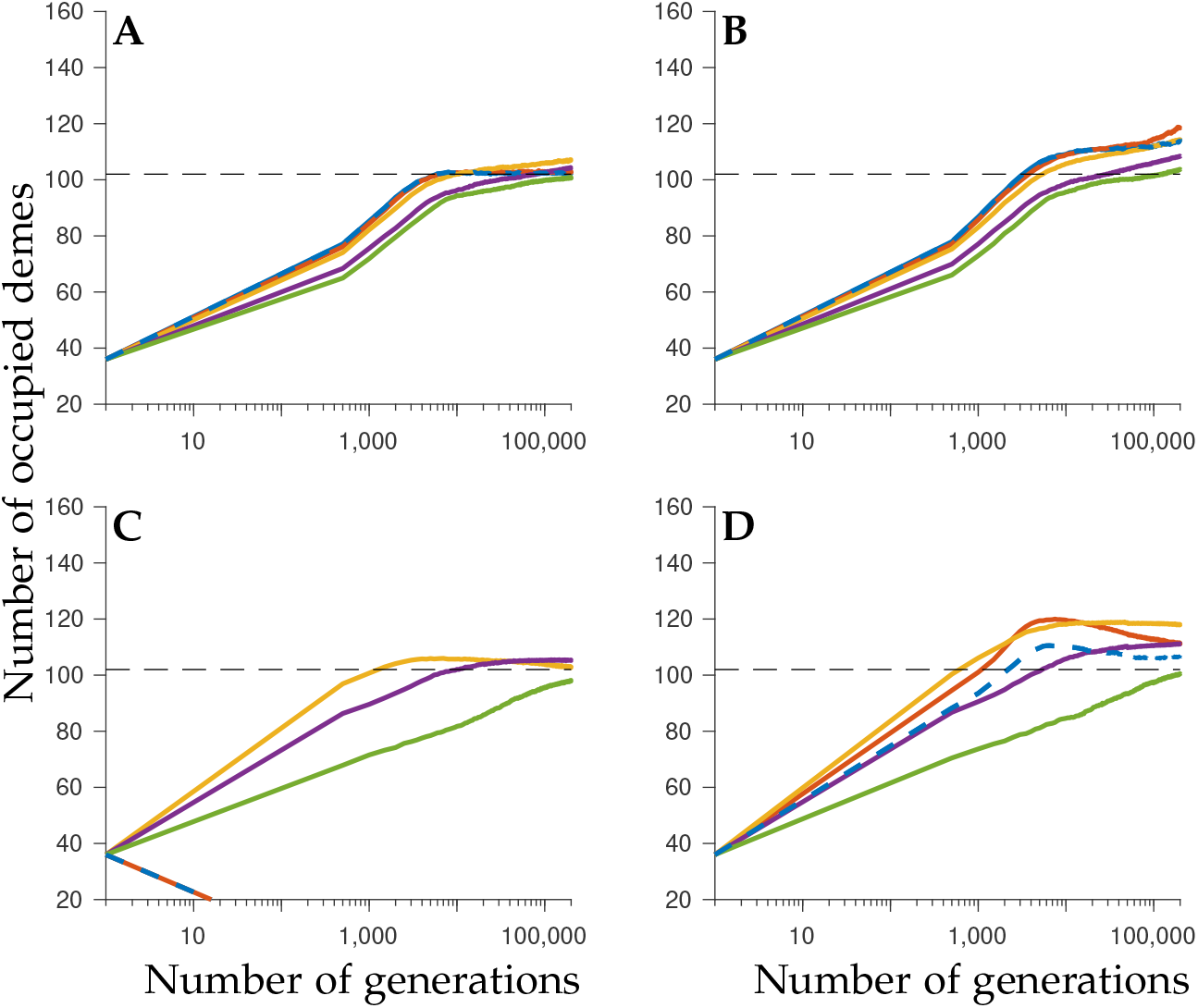
Temporal evolution of the mean number of occupied demes for strong selection. The results correspond to the model with low standing genetic variation (parameter set LS; A, B) or high standing genetic variation (parameter set HS; C, D). Selfing is either not allowed (A, C) or it is allowed at no cost (B, D). Dashed lines show the expected number of occupied demes in the steady state. Different colours denote the results with different recombination rates: *c* = 0.5 (i.e. free recombination; dashed blue), *c* = 5 ⋅ 10^*−*3^ (red), *c* = 10^*−*3^ (yellow), *c* = 10^*−*4^ (purple), *c* = 10^*−*5^ (green).

Recall that we ran the simulations for 200, 000 generations after the start of the expansion. When selfing was not allowed, and the population was initialised with low standing genetic variation, the total expansion range approximately attained the steady state within this time period (figs. 1A–2A). In these cases, the expansion range in the steady state agreed well with the expected range predicted by Polechová and Barton (2015) (see dashed lines in figs. 1A–2A, and Fig. 3A, E). The same was true for the model with high standing genetic variation, no selfing and strong selection when recombination was not too low, i.e. *c* > 10^*−*5^ (figs. 2C, 3G). In other cases, however, the steady state was not reached during the simulated 200, 000 generations (figs. 1B-D, 2B,D). Importantly, the dynamics of range expansion towards the steady state differed critically between the models initialised with low and high standing variation. In the model initialised with low standing genetic variation, the range increased approximately monotonically, and, in some cases, it stabilised (see above). However, when the simulations were initialised with high standing genetic variation, the range rapidly expanded reaching a maximum within 10, 000 generations after the start of the expansion when *c* ≥ 10^*−*3^ (or after approximately 100, 000 generations when *c* = 10^*−*4^), after which the range started to contract (figs. 1C-D, 2C-D). In many cases, the range contracted below the critical range (fig. 1C-D) when the range expansion was driven by standing genetic variation rather than novel mutations. When the mutation rate was increased from 10^*−*8^ to 10^*−*6^, however, the range contraction stopped at the critical range (fig. A2). The maximum range attained was larger than the expected critical range for freely recombining loci (denoted by dashed lines in figs. 1–2). Reduced recombination (10^*−*4^ ≤ *c* ≤ 10^*−*2^) significantly increased the persistence time beyond the critical range (table A2, table A3), while the maximal range was still as large as, or larger than, the maximal range obtained when *c* = 0.5, unless *c* was approximately 10^*−*4^ or smaller (see fig. 4; table A4-A5). Allowance of selfing typically increased the time to reach the maximum range (cf. A and B or C and D in fig. 4) and significantly increased the persistence time above the critical range for larger values of recombination rate, i.e. when *c* ≥ 10^*−*3^ (table A6). Selfing decreased the maximal range when *c* ≤ 5 ⋅ 10^*−*3^ and increased the maximal range when *c* ≥ 10^*−*3^ (table A7).

**Figure 3:**
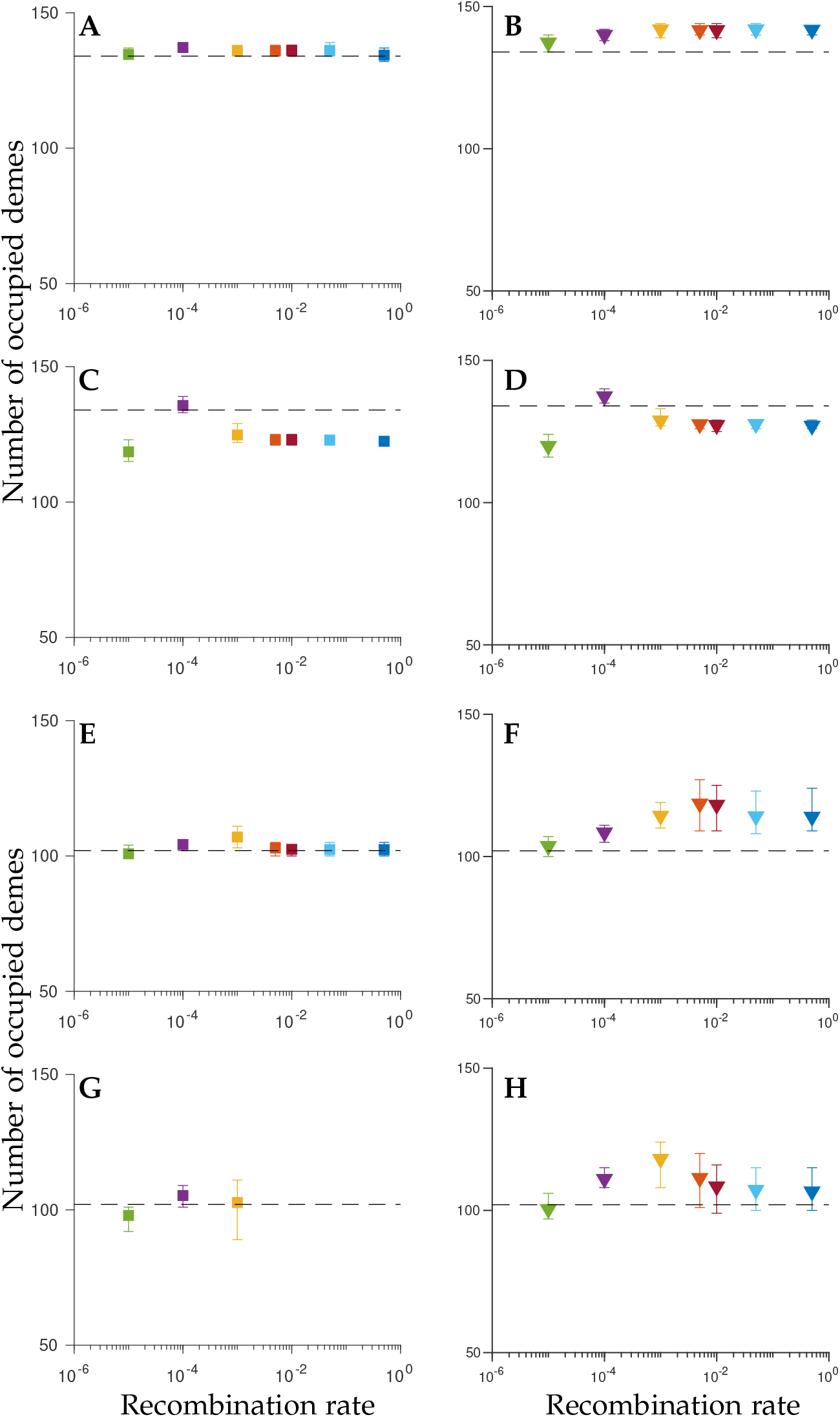
The number of occupied demes 200, 000 generations after the start of the expansion for the four parameter sets considered: parameter set LW (A, B), parameter set LS (C, D), parameter set HW (E, F) parameter set HS (G, H). Selfing is either not allowed (A, C, E, G) or it is allowed at no cost (B, D, F, H). Dashed lines show the expected number of occupied demes in the steady state. Different colours denote the results with different recombination rates: *c* = 0.5 (blue), *c* = 5 ⋅ 10^*−*3^ (red), *c* = 10^*−*3^ (yellow), *c* = 10^*−*4^ (purple), *c* = 10^*−*5^ (green). The bars indicate the 2.5 and 97.5 percentiles of 50 realisations.

**Figure 4:**
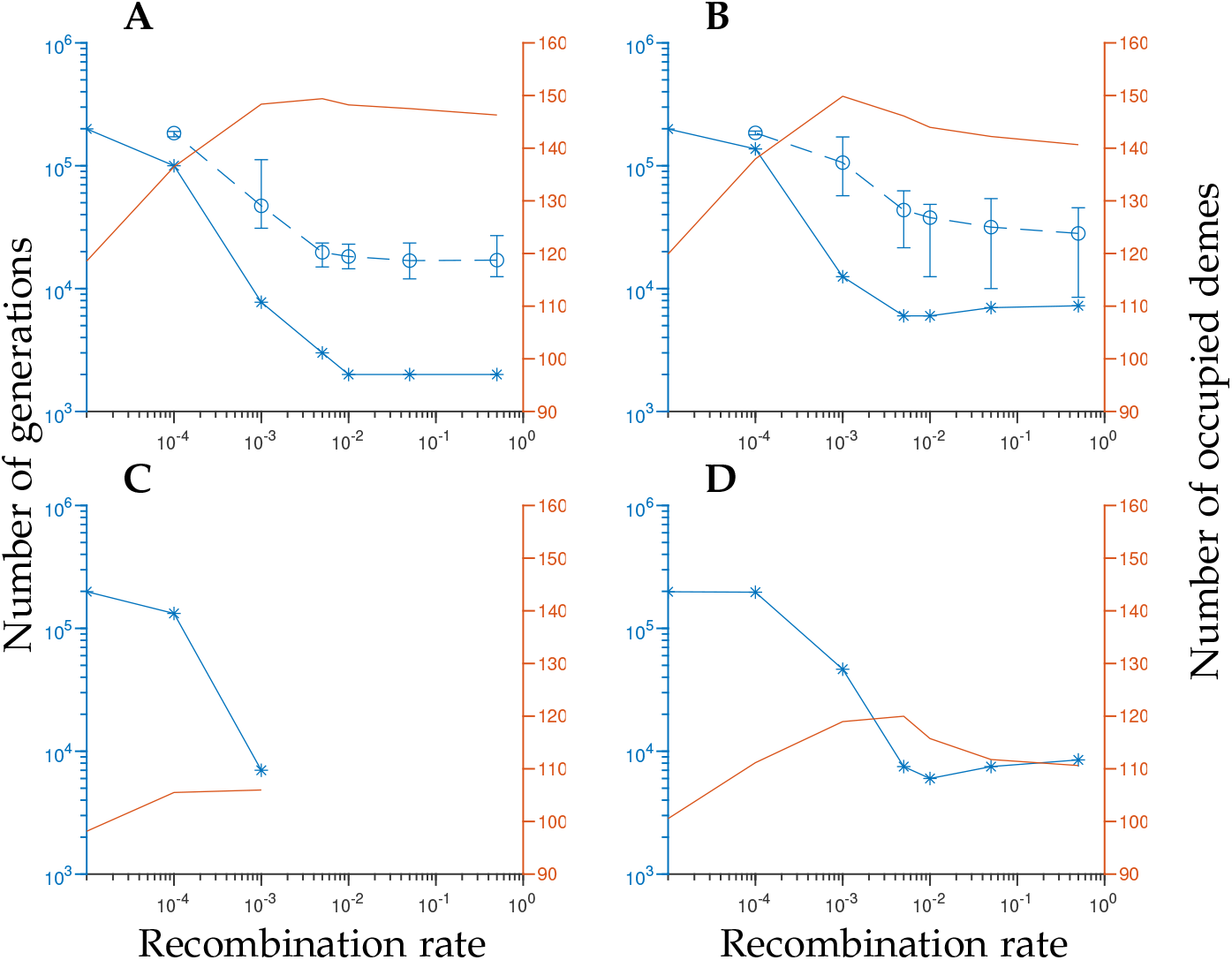
Average time to reach the maximal range extent (blue, left vertical axis, stars), average persistence time above the critical range (blue, left vertical axis, rings) and average maximal range extent (orange, right vertical axis, solid lines) when simulations were initialised with high genetic variation. Bars correspond to the minimum and maximum persistence time observed. The results correspond to the model with weak selection per locus (parameter set HW; A, B) or strong selection per locus (parameter set HS; C, D). Selfing is either not allowed (A, C) or it is allowed at no cost (B, D). The persistence time is not shown for parameter set HS, because the range of the population either did not contract below the critical range once it had exceeded it, or the population never reached the critical range.

A reduction of the recombination rate between the adaptive loci (but not below *c* = 10^*−*4^) led to a statistically significant increase of the total range occupied 200, 000 generations after the start of the expansion when selfing was not allowed, compared to when there was free recombination between the adaptive loci and selfing was not allowed (table A8). However, when the recombination rate was reduced and selfing was allowed there were no obvious trends, although some reduced recombination rates led to a significantly increased range (table A9). Likewise, the allowance of selfing led to a statistically significant increase of the total range 200, 000 generations after the start of the expansion compared to when selfing was not allowed, for each recombination rate and each parameter set tested (table A10). However, due to relatively large stochastic variations between different realisations of the model only some realisation with reduced recombination or with selfing allowed led to a greater range than that attained with free recombination or without selfing, respectively (fig. 3).

## Discussion

Using individual-based simulations, we have demonstrated that the genetic architecture of adaptive loci, as well as the mode of reproduction and the amount of standing genetic variation in the source population, are important factors during range expansions. In what follows we list and discuss our main findings.

First, we demonstrated that populations with reduced recombination rate between the adaptive loci may more efficiently utilise genetic variation for range expansions, compared to populations with free recombination. Recall that, since populations may adapt to new environments either by new mutations or by standing genetic variation (Barret and Schluter 2008), we examined two extremes of the possible amounts of genetic variation. One is the minimum amount of genetic variation required for adaptation to the central part of the habitat, and a rather high mutation rate (*µ* = 10^*−*6^). The other is the maximum possible amount of genetic variation, but such that the trait mean of the population still tracks the optimal phenotype over the central habitat and the adaptive loci are in linkage equilibrium. We used a lower mutation rate (*µ* = 10^*−*8^) when the simulations were initialised with high genetic variation. The extremely high genetic load, imposed on the population by the latter starting condition, inevitably caused the growth rate during the first generation to be strongly negative in every simulation performed. This was followed by an increase in the mean fitness in the subsequent generations, as poorly adapted individuals were purged from the population. However, when selection per locus was strong, no selfing was allowed and there was free recombination rate between the adaptive loci, the mean fitness failed to increase sufficiently fast and the population went extinct after a few generations. By contrast, selfing and/or reduced recombination prevented extinction of the population, even when selection was strong and the initial genetic variation was high. We suggest two different explanations to why selfing or reduced recombination, respectively, could rescue a population with this amount of genetic variation from extinction. When selfing is allowed, those individuals that, by chance, happen to have high fitness can produce a large number of gametes and with a high probability mate with themselves at the expansion front, because of the small population size. This rapidly eliminates locally poorly adapted alleles, while well-adapted alleles are proliferated, and thus the population is rescued from extinction. When the recombination rate is reduced, there is a high probability that gametes do not undergo recombination, and hence an individual with at least one homologous set of favourable alleles may have an offspring of high fitness. By contrast, when recombination between the adaptive loci is free (*c* = 0.5) the fitness of the offspring will approach the average fitness of the population, and therefore the population will go extinct after a few generations, because the average fitness of the population fails to increase sufficiently fast.

An important difference between the outcome of simulations with selfing and with reduced recombination, is that selfing quickly reduces heterozygosity and genetic variation when mating occurs at random in small populations (such as at the expansion front) (Stebbins 1957; Wright et al. 2013), whereas reduced recombination does not have this effect. We note that our model includes dominance effects. Indeed, while the allelic effects in our model were additive for the trait, they were not additive for the fitness. Instead, the fitness depended on the interaction between alleles, including the effect of dominance. The consequence of this is most easily seen in the case when the initial genetic variation is high, the selection per locus is weak (and hence extinction did not occur in our simulations) and when *c* ≥ 5 ⋅ 10^*−*3^. Allowance of selfing under these conditions led to a slower range expansion and a smaller maximal range, when compared to the same conditions but without selfing, because when the genetic variation is high, selfing increases the chances of producing homozygotes for a potentially recessive, locally deleterious allele.

Second, we found that reduced recombination between the adaptive loci decreases the speed of range expansion over heterogeneous environments. Despite this, however, we demonstrated that reduced recombination may increase the maximum range attained by the population. Indeed, slower range expansion leads to weaker bottlenecks (Gilbert et al. 2017, 2018), which makes local selection more efficient relative to drift. This suggests that there is a self-regulating mechanism built in for range expansions over heterogeneous environments: when a reduction in the recombination rate between the adaptive loci leads to an increase of the genetic load at the expansion front, the range expansion slows down, which in turn prevents the genetic load from building up too fast. Thus, the effects of increased genetic load due to reduced recombination, and decreased drift due to slower range expansion partly cancel each other out. At the same time, a reduction in the recombination rate preserves the genetic variation present in the population, because blocks of closely linked alleles are essentially cloned from a generation to the next. These blocks of alleles may be beneficial, and thus act as a single locus with a large beneficial allele-effect, and hence be under stronger selection than single alleles. A similar effect was observed by Polechová and Barton (2015) when the allele-effect sizes were randomly (exponentially) distributed. The range margins eventually became dominated by large-effect alleles which were under stronger selection the the average allele, and hence the range could expand a few demes longer than expected from the mean selection per locus. As a consequence of the combined effects of well-adapted blocks of alleles and a more moderate effect of drift than one might expect, the maximal range extent may be larger when the recombination rate is reduced compared to when the recombination is free. This applies both to range expansions driven by standing genetic variation, and range expansions driven by novel mutations (in the latter case, the maximum range equals the range attained at the end of our simulations).

Third, we showed that reduced recombination may considerably increase the time that the population spends beyond the critical range, when the standing genetic variation was high, the selection per locus was weak and selfing was not allowed in our simulations. This is because tighter linkage more efficiently protects blocks of locally well-adapted alleles. This is akin to the mechanism discussed by Kirkpatrick and Barrett (2015) *albeit* under a simpler model with locally beneficial alleles harboured by a chromosomal inversion. Interestingly, when our simulations were initialised with high standing genetic variation, and mutation rate was low, we noticed that the range contracted to a smaller range than the critical range. This finding highlights a difference between range expansions due to novel mutations and due to standing genetic variation. When range expansion is driven by novel mutations, the range increases monotonically to (or slightly beyond, see discussion in the previous paragraph) the critical range, whereas range expansions driven by standing genetic variation eventually halt whereupon the range contracts. In such cases, the range may contract to a smaller range than expected, because selection acts to eliminate excess variation and in the process may eliminate some of the background variation required for adaptation to the entire critical range. In these cases, a moderately reduced recombination rate increases the persistence time above the critical range, and when the recombination rate is further decreased (to *c* = 10^*−*4^ in our simulations) the range contraction may be completely prevented. This once again emphasises the importance of reduced recombination for maintenance of genetic variation.

Fourth, we found that selfing could increase the long-term range extent considerably, during range expansion over heterogeneous environments. This seems consistent with empirical evidence that selfing occurs relatively frequently in range margins (Pujol et al. 2009). Under constant environmental conditions, it has been suggested that selfing may be beneficial during range expansions because a single individual may be sufficient to establish a colony (Baker 1955; Tomlinson 1966; Pannell and Barrett 1998). We suggest that this is a part of the explanation to why selfing is advantageous in our case as well. Notably, our results also suggest that a colony originating from a single selfing individual, especially if this individual is (by chance) well adapted to the local habitat, may partly block gene flow from the source population (fig. A3, fig. A4) by forming a high density population with little or no genetic variation. Because of the large population size of individuals with near-optimal fitness, it would be highly unlikely for locally maladapted alleles originating from the source population to establish in this colony. A similar effect (*albeit* with clonal reproduction rather than selfing, and in a model with a homogeneous environment) was observed in a modelling study by Rafajlović et al. (2017) where wide spread long-lived clonal colonies were formed during range expansion of a partly clonal population. A well-known disadvantage of selfing is the increased frequency of individuals homozygous for recessive deleterious alleles (Charlesworth and Charlesworth 1987; Pujol et al. 2009). As mentioned above, this is captured by our model with high standing genetic variation and weak selection, wherein selfing populations expand their ranges slower than non-selfing ones. Further theoretical studies are required to establish how the allowance of selfing, with rates of selfing that are higher (fixed or evolving) than those used in our model, affects range expansion over heterogeneous environments when taking into account both dominance and genetic drift.

For an empirical example of how genetic architecture may be of importance to range expansions we mention the population of turbot (*Scophthalmus maximus*) that recently (within the last 8, 000 years) have colonised the Baltic Sea (Le Moan et al. 2019). In the Baltic Sea population of turbot the genomic regions where differences, compared to the marine populations, are concentrated have low recombination rates compared to the genomic average (Le Moan et al. 2019). Our results suggest a possible explanation to why reduced recombination could have benefited the Baltic Sea population of turbot. There is a short segment of rapidly decreasing salinity through the Kattegat and the Danish straits (Maar et al. 2011). The environmental gradient in this region might be too steep to cross for a populations with free recombination between the loci that are involved in tolerance to different salinity levels. Our results, however, suggest that reduced recombination may have been the key factor that helped the turbot population to cross this, otherwise insurmountable, barrier. Our results further suggest that many populations that inhabit the Baltic Sea, such as the turbot, may potentially undergo a range contraction in the future, if their range expansion to the Baltic Sea was primarily driven by standing genetic variation rather than novel mutations. To test this hypothesis, further empirical and theoretical work is required. While our study did not treat chromosomal inversions *per se*, they are closely related to tight genetic architectures. Chromosomal inversions have frequently been implicated in promoting range expansions. For example, this has been empirically observed in species such as threespine stickleback (*Gasteroseteus aculeatus*; Jones et al. 2012), yellow monkeyflower (*Mimulus guttatus*; Lowry and Willis 2010), rough periwinkle (*Littorina saxatilis*; Johannesson et al. 2017; Westram et al. 2018; Faria et al. 2019) and red imported fire ant (*Solenopsis invicta*; Tsutsui and Suarez 2003; Wang et al. 2013). Our results that are pertinent to reduced recombination in general, are likely to also be applicable to inversions. Similarly, uniparental reproduction, such as cloning (Ting and Geller 2000; Kearney 2003; Kliber and Eckert 2005; Tatarenkov 2005; Kawecki 2008) or selfing (Pujol et al. 2009), is relatively frequently occurring in populations that recently have undergone range expansions, or in populations that live in marginal environments. Our results suggest that selfing may be promoted in range margins because it may greatly reduce the negative impact of locally maladapted migrants, and that this effect may persist for a very long time, even when selfing mainly occurs during the initial stage of colonisation.

In conclusion, the results presented here advance our understanding of how tight versus loose genetic architectures and reproductive strategies (selfing allowed or not) can influence range expansions. We suggest that further theoretical work on range expansions should take into account the genetic architecture of adaptive loci.

## Acknowledgments

The authors would like to thank Roger K. Butlin and Kerstin Johannesson for comments on the manuscript and many other colleagues from the Centre for Marine Evolutionary Biology at the University of Gothenburg (CeMEB; www.cemeb.science.gu.se), and Jitka Polechová for valuable discussions on the topic. This work was funded by the Hasselblad Foundation Grant to Female Scientists awarded to MR, by a grant from the Swedish Research Council Formas to MR, and it was additionally supported by grants from Swedish Research Councils (Formas and VR) to the CeMEB. The simulations were performed on resources at Chalmers Centre for Computational Science and Engineering (C3SE) provided by the Swedish National Infrastructure for Computing (SNIC).

## Statement of authorship

MR conceived and designed the study. ME and MR designed the simulations. ME wrote the computer code for the simulations and performed the simulations. ME carried out the analyses of the results. ME and MR interpreted the results. ME and MR wrote the first draft of the manuscript and edited it. MR supervised the study.

