## Supplementary Information for "The effect of genetic architecture and selfing on the capacity of a population to expand its range"

### Appendix A: Supplementary Figures and Tables

In this appendix, we present and briefly explain additional figures of relevance to our results. Fig. A1 demonstrates that the expansion speed increases when the recombination rate between the adaptive loci increases up to $c \approx 5 \cdot 10^{-3}$; when $c$ is increased further, the speed saturates.

Fig. A2 demonstrates that when the simulations were run with weak selection per locus and were initialised with high standing genetic variation, a relatively high mutation rate ($\mu = 10^{-6}$) leads to a range extent that agrees well with the critical range. By contrast, a lower mutation rate ($\mu = 10^{-8}$) results in a smaller range extent than the critical range 200,000 generations after the start of the expansion.

Figs. A3–A5 illustrate that large, and essentially isolated, populations with low genetic variation (fig. A4 B) may spontaneously form at the edge of the population when selection per locus is strong, the population has low standing genetic variation but high mutation rate ($\mu = 10^{-6}$), and selfing is allowed. Under the same conditions but with high standing genetic variation and lower mutation rate ($\mu = 10^{-8}$), isolated populations may remain above the critical range after the connected range has contracted following the initial range expansion (fig. A5). The genetic variation between these isolated populations may be surprisingly high because two very distinct populations come into contact here.

Fig. A6 shows that smaller recombination rate between the adaptive loci may preserve a higher amount of genetic variation for a longer time, compared to when the recombination rate is higher (compare panels A-D for low standing variation, and panels E-H for high standing genetic variation).

In table A1 it is shown that, when populations with reduced recombination are compared to populations with free recombination between the adaptive loci, the increase in the effective environmental gradient at the range margin at the end of the simulations (denoted by $B_{\text{end}}$) was in some cases large. When the recombination rate was very small ($c = 10^{-5}$) a smaller range extent than for free recombination, and hence a smaller effective environmental gradient at the expansion front, was reached 200,000 generations after the start of the expansion, due to the slow range expansion (hence the negative value in table A1).

For further results and their interpretation see Results, and Discussion in the main text.

Table A1: Increase in % of $B_{\text{end}}$ when the recombination rate between the adaptive loci was reduced.

| Recombination rate | 0.05 | 0.01 | $5 \cdot 10^{3}$ | $10^{-3}$ | $10^{-4}$ | $10^{-5}$ |
|---|---|---|---|---|---|---|
| Parameter set LW | 0% | 0% | +1% | +4% | +9% | +1% |
| Parameter set LS | +1% | +1% | +6% | +38% | +12% | −8% |

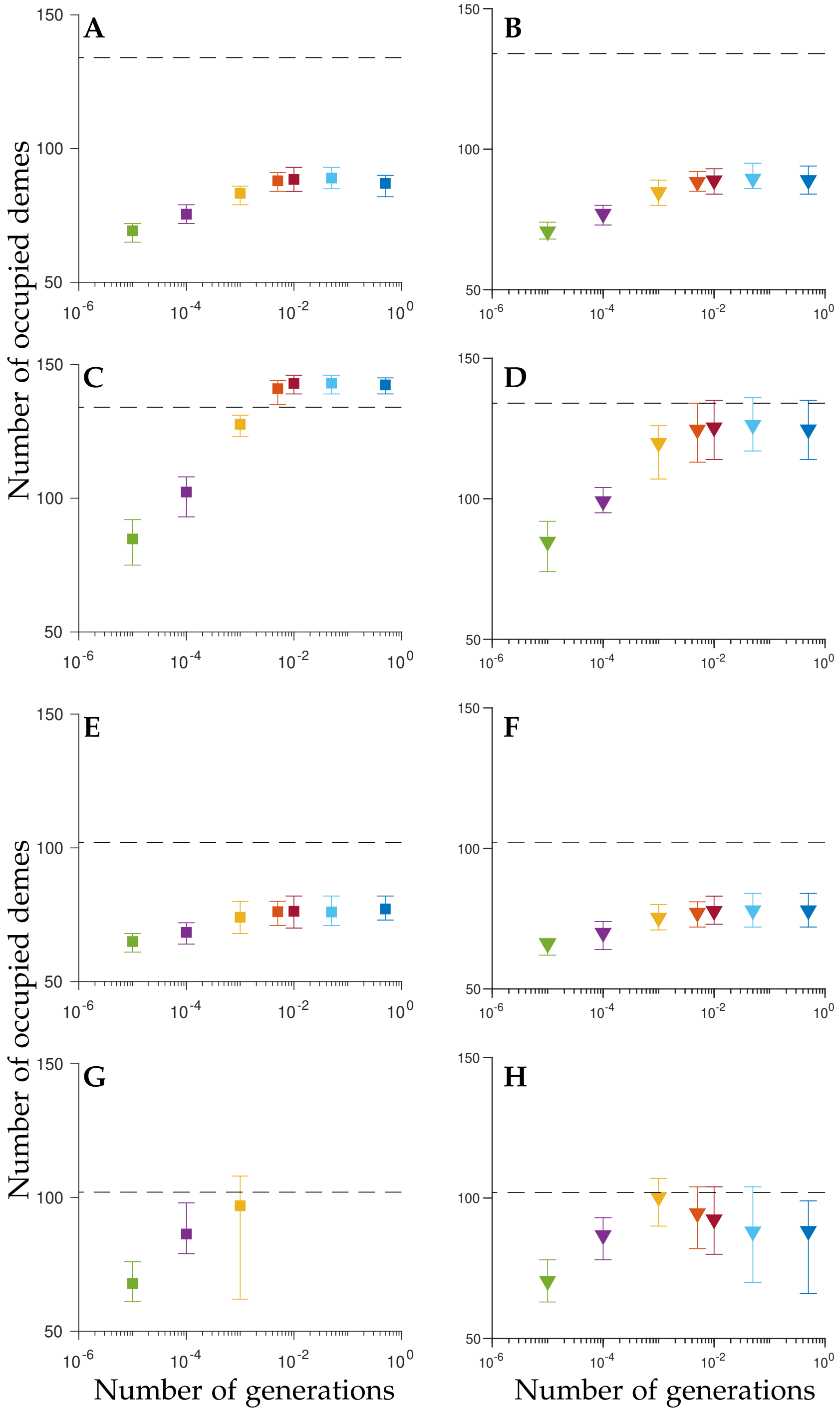


Figure A1: The number of occupied demes 500 generations after the start of the expansion for the four parameter sets considered: parameter set LW (A, B), parameter set LS (C, D), parameter set HW (E, F) parameter set HS (G, H). Selfing is either not allowed (A, C, E, G) or it is allowed at no cost (B, D, F, H). Dashed lines show the expected number of occupied demes in the steady state. Different colours denote the results with different recombination rates: $c = 0.5$ (blue), $c = 5 \cdot 10^{-3}$ (red), $c = 10^{-3}$ (yellow), $c = 10^{-4}$ (purple), $c = 10^{-5}$ (green). The bars indicate the 2.5 and 97.5 percentiles.

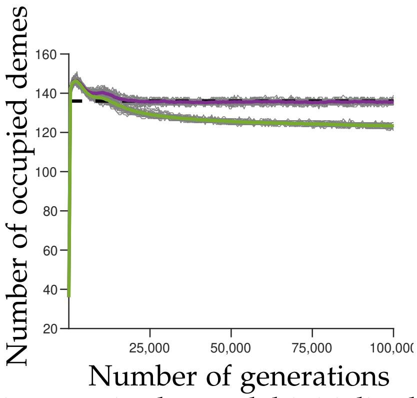


Figure A2: The effect of mutation rate in the model initialised with high standing genetic variation. The dashed line indicates the critical range. The purple line is the average number of occupied demes in the model with $\mu = 10^{-6}$, the green line is the average number of occupied demes in the model with $\mu = 10^{-8}$ (50 realisations in total). Grey lines show each individual realisation. Other parameters correspond to the parameter set HW, with $c = 0.5$ and no selfing allowed.

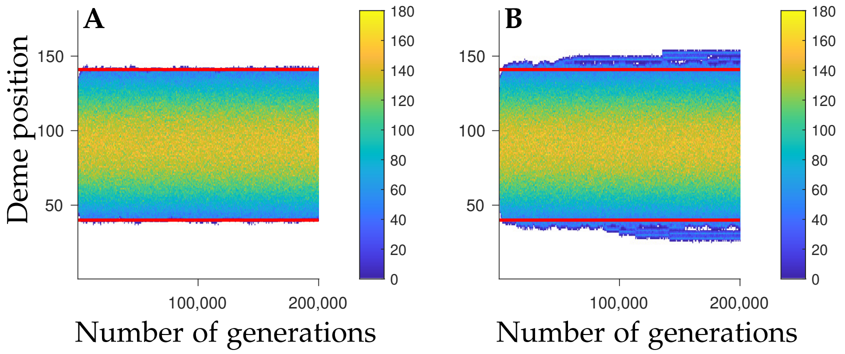


Figure A3: Evolution of local population size for single randomly chosen realisations with free recombination for parameter set LS (low initial standing genetic variation). Selfing is either not allowed (A) or it is allowed at no cost (B). Horizontal red lines indicate the critical range, white regions denote empty demes.

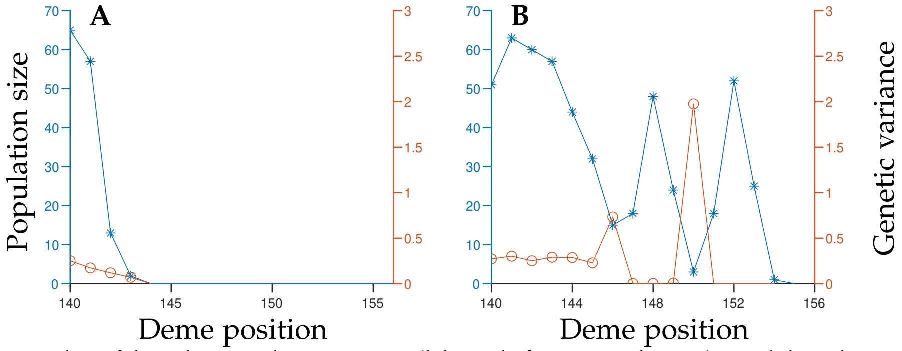


Figure A4: Example of local population size (blue, left vertical axis) and local genetic variance (orange, right vertical axis), at the edge of the range distribution between deme 140 and 156, measured 200,000 generations after the start of the expansion. The data corresponds to figure A3.

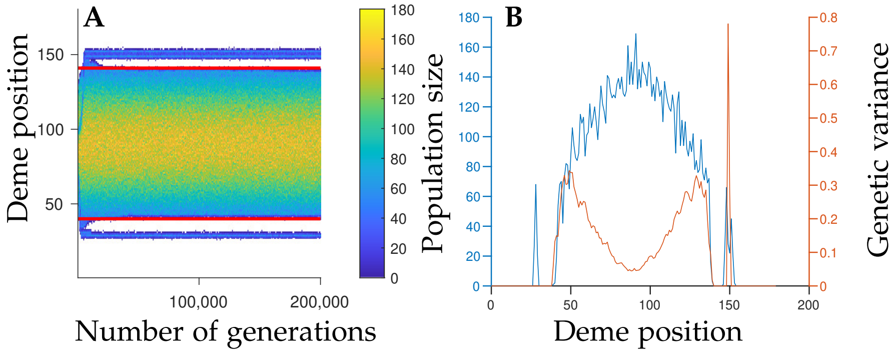


Figure A5: A: Evolution of local population size for single randomly chosen realisations with free recombination for parameter set HS (high initial standing genetic variation) with selfing allowed. Horizontal red lines indicate the critical range, white regions denote empty demes. B: Cross section of the local population size (blue, left vertical axis) and local genetic variance (orange, right vertical axis) 200, 000 generations after the start of the expansion.

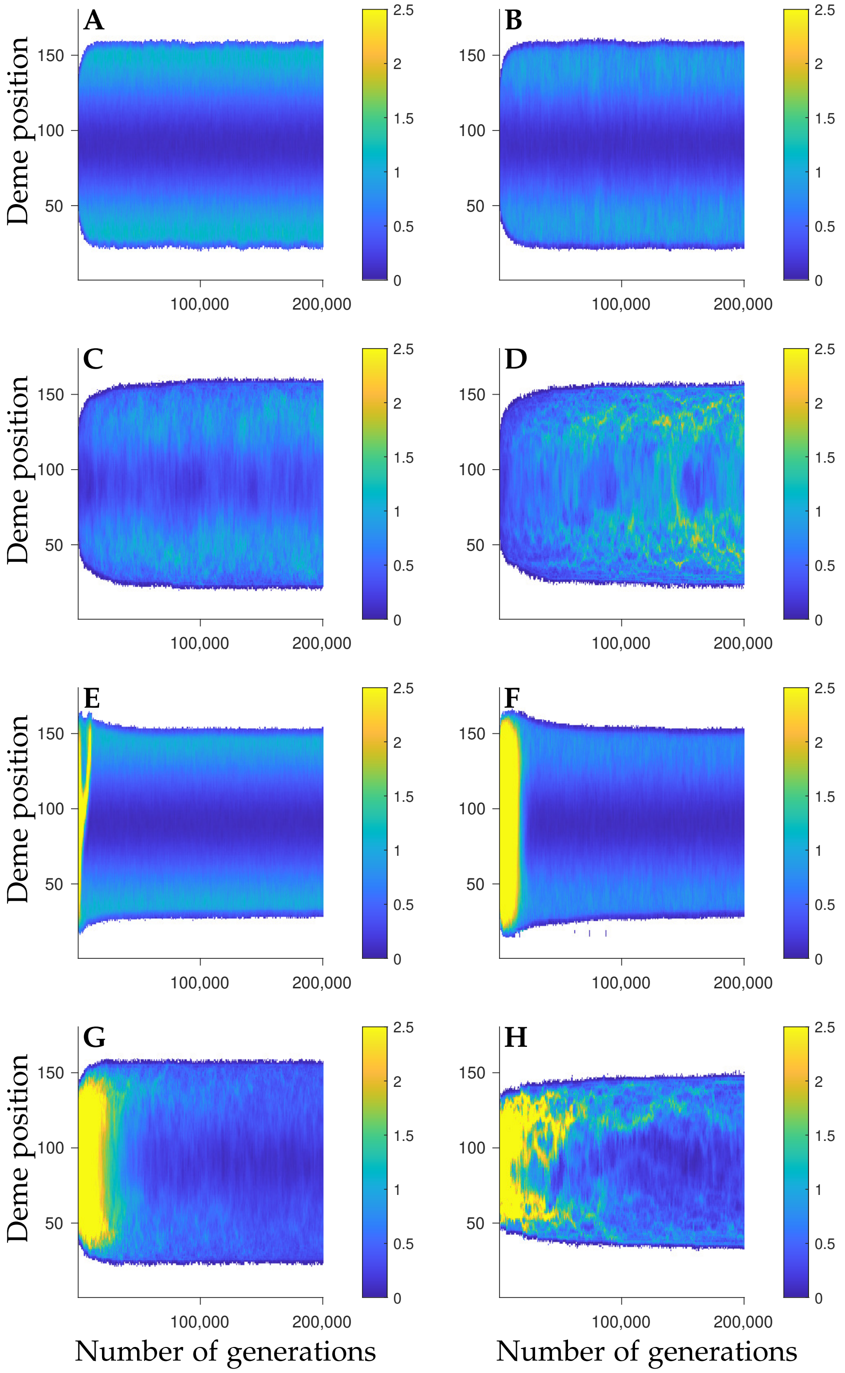


Figure A6: Evolution of local genetic variation for single randomly chosen realisations with different recombination rates for weak selection and without selfing. The results correspond to free recombination (A, E), $c = 10^{-3}$ (B, F), $c = 10^{-4}$, (C, G) and $c = 10^{-5}$ (D, H). The standing genetic variation is either low (A-D), or high (E-H). To make comparison between the high and low initial variation case easier, the values of $V_G$ were truncated to the range between 0 and 2.5. The maximum genetic variance obtained in E-H was about $V_G = 11$.

Next, we present tables of $p$-values.

Tables A2–A3 show the $p$-values obtained when the Wilcoxon-Mann-Whitney test was applied to the persistence time above the critical range for the simulations where the range expanded beyond the critical range and then contracted below it, namely when the selection per locus was weak and the initial standing genetic variation was high. We tested if reduced recombination increases the persistence time compared to when $c = 0.5$ when selfing was not allowed (table A2) and when selfing was allowed at no cost (table A3).

Tables A4–A5 show the $p$-values obtained when the Wilcoxon-Mann-Whitney test was applied to test whether the maximal range attained under the same conditions as in tables A2–A3 was increased when recombination was reduced. The $p$-values are shown both when selfing was not allowed (table A4), and when selfing was allowed at no cost (table A5).

In table A6 we present the $p$-values obtained when the Wilcoxon-Mann-Whitney was applied to test whether the persistence time increased when selfing was allowed, compared to when selfing was not allowed. Table A7, shows the $p$-values obtained when the Wilcoxon-Mann-Whitney test was applied both to test whether the maximal range was increased and whether the maximal range was decreased when selfing was allowed, compared to when selfing was not allowed. The tests were performed for each recombination rate considered in our study.

Finally, we present the $p$-values obtained when the Wilcoxon-Mann-Whitney test was applied to test whether the distribution of the total expansion ranges $200,000$ generations after the start of the expansion was increased when the recombination rate was smaller than $c = 0.5$. This was done both when selfing was not allowed (table A8) and when selfing was allowed at no cost (table A9). In table A8 no $p$-values are shown for parameter set HS, because all populations went extinct when the adaptive loci were recombining freely. When the recombination rate was very low ($c \leq 10^{-5}$ in table A8 or $c \leq 10^{-4}$ in table A9), the range expansion was very slow. Therefore, the null hypothesis for these cases cannot be rejected for the the time span we simulated here.

Table A10 shows the $p$-values obtained when the Wilcoxon-Mann-Whitney test was applied to test whether allowance of selfing led to a larger range compared to when selfing was not allowed for each recombination rate considered in the study.

Table A2: List of $p$-values for the null hypothesis that reduced recombination rate does not increase the persistence time above the critical range, when selfing was not allowed.

| Recombination rate | $p$-value |
|---|---|
| 0.05 | 0.384 |
| 0.01 | $7.27 \cdot 10^{-4*}$ |
| 0.005 | $2.99 \cdot 10^{-8*}$ |
| 0.001 | $3.35 \cdot 10^{-18*}$ |
| $10^{-4}$ | $3.31 \cdot 10^{-18*}$ |
| $10^{-5}$ | 1.00 |

Note: The parameters correspond to parameter set HW: the number of adaptive loci was $L = 574$, the allele-effect size was $\alpha = \frac{1}{\sqrt{20}}$, the width of stabilising selection was $V_S = 2$, the mutation rate was $\mu = 10^{-8}$ and the initial standing genetic variation was high. The asterisk (*) denotes a significant $p$-value for which the null hypothesis can be rejected. The null hypothesis was that a reduction of recombination rate does not increase the persistence time relative to when $c = 0.5$. The alternative hypothesis was that the persistence is increased when $c$ is decreased.

Table A3: List of $p$-values for the null hypothesis that reduced recombination rate does not increase the persistence time above the critical range, when selfing was allowed.

| Recombination rate | $p$-value |
|---|---|
| 0.05 | 0.0633 |
| 0.01 | $4.79 \cdot 10^{-8}$* |
| 0.005 | $1.86 \cdot 10^{-12}$* |
| 0.001 | $3.51 \cdot 10^{-18}$* |
| $10^{-4}$ | $3.44 \cdot 10^{-18}$* |
| $10^{-5}$ | 1.00 |

Note: The parameters correspond to parameter set HW: the number of adaptive loci was $L = 574$, the allele-effect size was $\alpha = \frac{1}{\sqrt{20}}$, the width of stabilising selection was $V_S = 2$, the mutation rate was $\mu = 10^{-8}$ and the initial standing genetic variation was high. The asterisk (*) denotes a significant $p$-value for which the null hypothesis can be rejected. The null hypothesis was that a reduction of recombination rate does not increase the persistence time relative to when $c = 0.5$. The alternative hypothesis was that the persistence is increased when $c$ is decreased.

Table A4: List of $p$-values for the null hypothesis that reduced recombination rate does not increase the maximal range extent, when selfing was not allowed.

| Recombination rate | $p$-value |
|---|---|
| 0.05 | $5.62 \cdot 10^{-5}$* |
| 0.01 | $2.70 \cdot 10^{-13}$* |
| 0.005 | $1.00 \cdot 10^{-16}$* |
| 0.001 | $2.44 \cdot 10^{-14}$* |
| $10^{-4}$ | 1.00 |
| $10^{-5}$ | 1.00 |

Note: The parameters correspond to parameter set HW: the number of adaptive loci was $L = 574$, the allele-effect size was $\alpha = \frac{1}{\sqrt{20}}$, the width of stabilising selection was $V_S = 2$, the mutation rate was $\mu = 10^{-8}$ and the initial standing genetic variation was high. The asterisk (*) denotes a significant $p$-value for which the null hypothesis can be rejected. The null hypothesis was that a reduction of recombination rate does not increase the maximal range relative to when $c = 0.5$. The alternative hypothesis was that the maximal range is increased when $c$ is decreased.

Table A5: List of $p$-values for the null hypothesis that reduced recombination rate does not increase the maximal range extent, when selfing was allowed.

| Recombination rate | $p$-value |
|---|---|
| 0.05 | $0.0052^{*}$ |
| 0.01 | $4.00 \cdot 10^{-7*}$ |
| 0.005 | $9.55 \cdot 10^{-13*}$ |
| 0.001 | $2.43 \cdot 10^{-17*}$ |
| $10^{-4}$ | 1.00 |
| $10^{-5}$ | 1.00 |

Note: The parameters correspond to parameter set HW: the number of adaptive loci was $L = 574$, the allele-effect size was $\alpha = \frac{1}{\sqrt{20}}$, the width of stabilising selection was $V_S = 2$, the mutation rate was $\mu = 10^{-8}$ and the initial standing genetic variation was high. The asterisk (*) denotes a significant $p$-value for which the null hypothesis can be rejected. The null hypothesis was that a reduction of recombination rate does not increase the maximal range relative to when $c = 0.5$. The alternative hypothesis was that the maximal range is increased when $c$ is decreased.

Table A6: List of $p$-values for the null hypothesis that allowance of selfing does not increase the persistence time above the critical range.

| Recombination rate | $p$-value |
|---|---|
| 0.5 | $1.25 \cdot 10^{-9*}$ |
| 0.05 | $1.25 \cdot 10^{-13*}$ |
| 0.01 | $6.43 \cdot 10^{-17*}$ |
| 0.005 | $5.56 \cdot 10^{-18*}$ |
| 0.001 | $2.96 \cdot 10^{-16*}$ |
| $10^{-4}$ | 0.301 |
| $10^{-5}$ | 1.00 |

Note: The parameters correspond to parameter set HW: the number of adaptive loci was $L = 574$, the allele-effect size was $\alpha = \frac{1}{\sqrt{20}}$, the width of stabilising selection was $V_S = 2$, the mutation rate was $\mu = 10^{-8}$ and the initial standing genetic variation was high. The asterisk (*) denotes a significant $p$-value for which the null hypothesis can be rejected. The null-hypothesis was that selfing does not increase the persistence time, for a given value of $c$, compared to when selfing is not allowed. The alternative hypothesis was that selfing does increase the persistence time for the given value of $c$.

Table A7: List of $p$-values for the null hypotheses that allowance of selfing does not change the maximal range extent.

| Recombination rate | $p$-value for decrease | $p$-value for increase |
|---|---|---|
| 0.5 | $1.30 \cdot 10^{-15*}$ | 1.00 |
| 0.05 | $1.77 \cdot 10^{-15*}$ | 1.00 |
| 0.01 | $1.68 \cdot 10^{-14*}$ | 1.00 |
| 0.005 | $2.59 \cdot 10^{-9*}$ | 1.00 |
| 0.001 | 1.00 | $1.00 \cdot 10^{-5*}$ |
| $10^{-4}$ | 1.00 | $7.81 \cdot 10^{-8*}$ |
| $10^{-5}$ | 1.00 | 0.0036* |

Note: The parameters correspond to parameter set HW: the number of adaptive loci was $L = 574$, the allele-effect size was $\alpha = \frac{1}{\sqrt{20}}$, the width of stabilising selection was $V_S = 2$, the mutation rate was $\mu = 10^{-8}$ and the initial standing genetic variation was high. The asterisk (*) denotes a significant $p$-value for which the null hypothesis can be rejected. The two null hypotheses were 1. allowance of selfing does not decrease the maximal range relative to when selfing is not allowed, and 2. allowance of selfing does not increase the maximal range relative to when selfing is not allowed. The alternative hypotheses were that the maximal range is decreased when selfing is allowed and that the maximal range is increased when selfing is allowed, respectively.

Table A8: List of $p$-values for the null hypothesis that reduced recombination does not increase the range 200,000 generations after the start of the expansion, when selfing was not allowed.

| Recombination rate | Parameter set LW | Parameter set LS | Parameter set HW | Parameter set HS |
|---|---|---|---|---|
| 0.05 | $1.7 \cdot 10^{-11*}$ | 0.37 | 0.0036* | – |
| 0.01 | $2.2 \cdot 10^{-11*}$ | 0.25 | 0.0012* | – |
| 0.005 | $4.8 \cdot 10^{-10*}$ | 0.0042* | 0.0017* | – |
| 0.001 | $1.5 \cdot 10^{-10*}$ | $4.2 \cdot 10^{-17*}$ | $1.4 \cdot 10^{-9*}$ | – |
| $10^{-4}$ | $1.8 \cdot 10^{-17*}$ | $2.6 \cdot 10^{-11*}$ | $1.4 \cdot 10^{-18*}$ | – |
| $10^{-5}$ | 0.21 | 1.00 | 1.00 | – |

Note: The asterisk (*) denotes a significant $p$-value for which the null hypothesis can be rejected. The null hypothesis was that a reduction of recombination rate does not increase the range relative to when $c = 0.5$. The alternative hypothesis was that the range is increased when $c$ is decreased.

Table A9: List of $p$-values for the null hypothesis that reduced recombination does not increase the range 200,000 generations after the start of the expansion, when selfing was allowed.

| Recombination rate | Parameter set LW | Parameter set LS | Parameter set HW | Parameter set HS |
|---|---|---|---|---|
| 0.05 | 0.0014* | 0.38 | $1.1 \cdot 10^{-4}$* | 0.21 |
| 0.01 | 0.018* | $2.1 \cdot 10^{-6}$* | 0.065 | 0.022* |
| 0.005 | 0.17 | $1.5 \cdot 10^{-7}$* | 0.0012* | $2.8 \cdot 10^{-7}$* |
| 0.001 | 0.25 | $5.4 \cdot 10^{-3}$* | 0.86 | $1.7 \cdot 10^{-13}$* |
| $10^{-4}$ | 1.00 | 1.00 | $1.3 \cdot 10^{-18}$* | 0.11 |
| $10^{-5}$ | 1.00 | 1.00 | 1.00 | 1.00 |

Note: The asterisk (*) denotes a significant $p$-value for which the null hypothesis can be rejected. The null hypothesis was that a reduction of recombination rate does not increase the range relative to when $c = 0.5$. The alternative hypothesis was that the range is increased when $c$ is decreased.

Table A10: List of $p$-values for the null hypothesis that allowance of selfing does not increase the range 200,000 generations after the start of the expansion.

| Recombination rate | Parameter set LW | Parameter set LS | Parameter set HW | Parameter set HS |
|---|---|---|---|---|
| 0.5 | $2.3 \cdot 10^{-18}$* | $4.8 \cdot 10^{-18}$* | $2.1 \cdot 10^{-18}$* | $3.2 \cdot 10^{-20}$* |
| 0.05 | $1.3 \cdot 10^{-18}$* | $2.0 \cdot 10^{-18}$* | $8.0 \cdot 10^{-19}$* | $1.5 \cdot 10^{-20}$* |
| 0.01 | $1.6 \cdot 10^{-18}$* | $2.3 \cdot 10^{-18}$* | $1.5 \cdot 10^{-18}$* | $1.6 \cdot 10^{-20}$* |
| 0.005 | $1.5 \cdot 10^{-18}$* | $2.2 \cdot 10^{-18}$* | $1.2 \cdot 10^{-18}$* | $1.6 \cdot 10^{-20}$* |
| 0.001 | $2.7 \cdot 10^{-18}$* | $1.6 \cdot 10^{-17}$* | $7.6 \cdot 10^{-17}$* | $2.7 \cdot 10^{-17}$* |
| $10^{-4}$ | $2.3 \cdot 10^{-17}$* | $2.1 \cdot 10^{-17}$* | $4.2 \cdot 10^{-10}$* | $1.3 \cdot 10^{-17}$* |
| $10^{-5}$ | $4.4 \cdot 10^{-15}$* | $1.0 \cdot 10^{-17}$* | $1.0 \cdot 10^{-17}$* | $1.0 \cdot 10^{-17}$* |

Note: The asterisk (*) denotes a significant $p$-value for which the null hypothesis can be rejected. The null-hypothesis was that selfing does not increase the range, for a given value of $c$, compared to when selfing is not allowed. The alternative hypothesis was that selfing does increase the range for this value of $c$.

### Appendix B: Additional Methods

In this appendix, we present additional details regarding the model that was used (see Methods in the main text for the full model).

The habitat we considered was modelled as a one-dimensional chain of $M$ demes of equal sizes. The distances between the demes were measured in units of the number of demes. We assumed an environmental gradient along the habitat, such that the optimal phenotype in deme $i = 1, \ldots, M$, denoted by $\theta_i$, was assumed to be a cubic polynomial of $i$

$$\theta_i = 0.000499i^3 - 0.0894i^2 + 4.1000i. \quad \text{(B1)}$$

To avoid edge effects, we chose the value of $M$ in such a way that the number of demes is at least ten times the expected width of a cline, denoted by $w$ below (Barton 2001):

$$w = 4\frac{\sigma\sqrt{V_S}}{\alpha} \quad \text{(B2)}$$

where $\sigma$ denotes the standard deviation of the distance between parent and offspring and $V_S$ denotes the width of the stabilising selection. As in Polechová and Barton (2015); Bridle et al. (2019) we used this equation as a guide for determining the starting conditions. We used this equation as a guide to obtain the optimal genotypes, even though it was derived under the assumption that the the loci underlying the trait under selection were unlinked and that the environmental gradient was linear. For the parameters considered in this study ($\sigma = \frac{1}{\sqrt{2}}$, $V_S \leq 2$, $\alpha \geq \frac{1}{\sqrt{20}}$; see table 2 in the main text), we have $M \geq 10w$ when $M = 180$, which, throughout, was used as the number of demes in the model. The predicted cline widths with our parmeters are 17.9 and 8.0. The corresponding mean cline widths (calculated using $w_{pq} = 4\int_{-\infty}^{\infty} pq\mathrm{d}x$, as in Polechová and Barton (2011)) $200,000$ generations after the start of the expansion were close to these values at the centre of the habitat, but narrower closer to the range margins due to stronger drift, as expected according to Polechová and Barton 2011 (table B1, table B2). The average cline widths in the tables are calculated for simulations with free recombination between the adaptive loci. For reduced recombination rate, clines in frequencies had not formed at all loci, but for clinal loci the cline widths were similar to those in tables B1–B2 (not shown).

Table B1: Cline widths in the model with free recombination and without selfing.

| Deme position | Parameter set LW | Parameter set LS | Parameter set HW | Parameter set HS |
|---|---|---|---|---|
| 70-110 | 18.06 | 8.04 | 17.87 | – |
| 50-70 & 110-130 | 16.08 | 6.47 | 15.91 | – |
| 0-50 & 130-180 | 11.32 | 4.99 | 11.73 | – |

Note: Average cline width for clines with centres at deme positions in the ranges indicated in the left column.

Table B2: Cline widths in the model with free recombination and selfing.

| Deme position | Parameter set LW | Parameter set LS | Parameter set HW | Parameter set HS |
|---|---|---|---|---|
| 70-110 | 18.04 | 8.04 | 17.83 | 7.98 |
| 50-70 & 110-130 | 15.73 | 6.47 | 16.16 | 6.44 |
| 0-50 & 130-180 | 11.51 | 4.58 | 12.29 | 4.90 |

Note: Average cline width for clines with centres at deme positions in the ranges indicated in the left column.

To model migration within the habitat, we assumed Gaussian dispersal. We implemented Gaussian dispersal in our discrete habitat as follows. The probability for an individual to migrate from deme $i$ to deme $j$ ($j \neq 1, M$) was given by

$$m_{j,i} = \frac{1}{\sqrt{2\pi\sigma^2}} \int_{j-i-1/2}^{j-i+1/2} \mathrm{e}^{-\frac{t^2}{2\sigma^2}} \mathrm{d}t. \tag{B3}$$

Conversely, for migration to the edge demes (i.e. for $j = 1$ or $j = M$) the probabilites were given by

$$m_{1,i} = \frac{1}{\sqrt{2\pi\sigma^2}} \int_{-\infty}^{1-i+1/2} \mathrm{e}^{-\frac{t^2}{2\sigma^2}} \mathrm{d}t, \tag{B4}$$

and

$$m_{M,i} = \frac{1}{\sqrt{2\pi\sigma^2}} \int_{M-i-1/2}^{\infty} \mathrm{e}^{-\frac{t^2}{2\sigma^2}} \mathrm{d}t. \tag{B5}$$

Note that migration to the edge demes was likely rarely (if ever) realised because the farthest deme occupied by the populations in our model was more than 10 demes away from any of the habitat edges.

Further details regarding the model are given in Methods in the main text.
